# Within-flock differences in exploratory tendency and flock performance in a highly gregarious bird

**DOI:** 10.1101/2022.10.25.513662

**Authors:** Claudio Carere, Celine Audebrand, Florian Desigaux, Rianne Pinxten, Marcel Eens, Heiko G. Rödel, Patrizia d’Ettorre

## Abstract

How individual differences translate into group outcomes is a timely and debated issue. Recent studies, especially in social arthropods and fish, focus on diversity of personality traits. These studies suggest that the phenotypic group assortment by personality type of an animal group, including the presence of “keystone” individuals, leads to group-level personality differences and can strongly impact both group and individual outcomes. However, little attention has been given to the variation of a given trait within a group. Theory predicts that phenotypic homogeneity rather than heterogeneity yields the optimal group performance, especially in an anti-predatory context, but the experimental support includes mainly morphological traits, e.g. body size or colouration. Here, we focus on personality and group level differences in a highly gregarious bird, the European starling (*Sturnus vulgaris*). We investigated how different degrees of within-flock variation in exploration affect flock outcomes in exploratory behaviour and in escape response after a frightening stimulus. First, we established consistent individual differences in exploration. Then, flocks of 4 birds were formed to obtain gradual differences in mean and in variation of exploration scores among flocks. Flocks underwent an exploration test and a perturbation test. More exploratory individuals entered the test arena earlier, tended to start flying more rapidly and also stopped flying sooner upon frightening. Flocks with a more homogeneous distribution with respect to this personality trait were significantly faster to enter the test room, but no effect on the escape response emerged. The mean exploration tendency of the flock or the maximum exploration tendency of purported ‘key’ individuals within groups did not play a notable role in explaining such differences in group performance. Our results indicate that it is not the individual that predicts/drives the flock outcome, but rather a group feature, namely phenotypic variation within the group.

## INTRODUCTION

Many organisms, including humans, regularly group together and perform collective actions such as birds forming coordinated flocks to evade predators, forage or roost (Krause & Ruxton, 2002; Storms et al., 2019). However, individual members composing animal groups might vary in their phenotype. This within-group variability can vary itself among groups, which will therefore differ in having more or less phenotypic variance depending on the distribution of the different individuals in a given group (Jolles et al., 2020). Indeed, group-level differences are widespread in the animal kingdom (e.g., ant colonies, Bengston & Dornhaus, 2014; bird flocks, Carere et al., 2009; fish shoals, Jolles et al., 2018; honeybee colonies, Wray et al., 2011).

Recent studies in several animal species (see below) suggest that morphological, physiological, or behavioural traits play a role in group assortment, while aspects of group composition such as the mean, range, or variance in such phenotypic traits affect group outcomes and individual fitness with potential evolutionary implications (Farine et al., 2015; Killen et al., 2017; Sih & Watters, 2005). As stressed by recent reviews, there is an urgent need of empirical work across taxa to get a better picture of such effects and their implications (Farine et al., 2015; Jolles et al., 2020).

Consistent individual differences in behaviour (i.e., personality) are ubiquitous and have important ecological and evolutionary implications (Carere & Maestripieri, 2013; Laskowski et al., 2022; Réale et al., 2007; Wolf & Weissing, 2012). Personality differences are found in many social species and at the same time similar differences are found at the group level (e.g., Bengston & Dornhaus, 2014; Jolles et al., 2018; Wray et al., 2011). It is therefore predicted that personality differences at the individual level give rise to personality differences at the group level exerting important influences on group behaviour and functioning (Webster & Ward, 2011; Wolf & Krause, 2014). A current major line of investigation concerns this relationship between individual and group personality. Several empirical studies show that personality differences affect the group as a whole. For instance, groups composed by half shy and half bold fish were faster to explore a novel environment and to sample a foraging patch than groups composed by only shy or only bold fish (Dyer et al., 2009). By contrast, highly inactive individuals could negatively inhibit the group (Brown & Laland, 2002). In another fish study, no correlation emerged between group exploratory behaviour and group mean personality scores, but a minority of individuals with certain personality types influenced substantially group behaviour, as group exploration was delayed by the least active individuals and enhanced by the most social individuals (Brown & Irving, 2014). In an ant species, within-colony variation in aggression positively correlated with worker productivity (Modlmeier & Foitzik, 2011) and personality traits have been shown to affect fitness in several other ant species (Blight et al., 2016; Bengston et al., 2017; Maák et al., 2021). In general, individual personalities influence the collective behaviour in ants (Kolay et al., 2020).

Despite the growing number of studies on this topic, the general effects of the composition of personality types within a group are still not well understood. This may be due to the complexity of the dynamic interactions from the individuals to the group and vice versa, and also to the fact that many experiments with artificial composition of groups tend to exclude “moderate” personality types. As a consequence, the difference between personality scores within a group, i.e. its phenotypic homogeneity/heterogeneity as a group feature, is often overlooked. Phenotypic homogeneity is actually central in the study of group living and stems from evolutionary drivers such as predation. Theory predicts that individuals within groups should be phenotypically more similar than between groups because this would confer a more efficient anti-predatory performance with important consequences on grouping preference by assortment of individuals (Cattelan & Griggio, 2018; Krause et al., 1996; Krause & Ruxton, 2002; Ranta et al., 1994). Empirical studies, mostly in fish, carried out both in laboratory and field conditions confirmed the main prediction, yet they focused on morphological traits such as body size or pigmentation (e.g., Cattelan & Griggio, 2018; Hoare et al., 2000). To our knowledge, despite the vast body of literature on animal personality, no study considering personality traits has been carried out so far specifically tackling the issue of within-group phenotypic homogeneity of these traits among individual members of a group.

The majority of studies on individual versus group-level personality have been carried out in social arthropods (e.g., Carere et al., 2018; Keiser & Pruitt, 2014; Modlmeier & Foitzik, 2011), fish (e.g., Brown & Irving, 2014; Dyer et al., 2009; Jolles et al., 2017) and to a lesser extent in primates (e.g., Koski & Burkart, 2015), but still very few studies so far have considered birds that display commonly flocking behaviour in many ecological contexts, including predation. Field studies on social networks in great tits (*Parus major*) showed assortative flocking in relation to personality as well as within-group variation in personality, conferring most effective coordinated actions while foraging (Aplin et al., 2013, 2014).

Here, we focus on how consistent behavioural differences in exploration tendency affect group level outcomes in a highly gregarious bird. Exploratory tendency is a key behavioural trait commonly used to phenotype inter-individual differences in vertebrates (e.g., Bisconti et al., 2022; Dingemanse et al., 2002; Réale et al., 2007; Rödel et al., 2015). In particular, we examined how different degrees of within-flock variation in personality traits affect flock performances in an exploration task and in an escape response upon a frightening stimulus. We chose to test this in captive flocks of the European starling (*Sturnus vulgaris*). Starlings are widely distributed and show a remarkable collective behaviour based on self-organization as well as consistent personality traits at the individual level (Minderman et al., 2009; Thys et al., 2017a, b). Their groups are characterised by fission-fusion dynamics with frequent changes in group composition, which are particularly evident in an anti-predatory context (e.g., Carere et al., 2009; Hildebrandt et al., 2010; Feare, 1984; Goodenough et al., 2017; Storms et al., 2019).

We first characterized individuals for exploratory behaviour, including testing its short- and long-term repeatability over time. Then we constructed same-sized flocks in such a way that these showed gradual differences in mean exploration scores as well as variation of exploration scores within flocks. The behaviour of each individual was scored both when performing alone and when acting in the flock.

The central hypothesis of the study was that within-flock homogeneity in a personality trait, rather than the group mean, would affect flock-level performance. More specifically, we tested whether (*i*) low intra-flock variation in exploratory behaviour decreases latency of exploratory behaviour at the level of the entire flock; (*ii*) low intra-flock variation in exploratory behaviour affects the flock escape response.

## METHODS

### Ethics note

The experiments carried in this study complied with ethical guidelines of the University of Antwerp and Flemish and European laws regarding animal welfare, and adhere to the ASAB/ABS guidelines for the use of animals in behavioural research and teaching. Permission to capture starlings from the wild and house them in captivity (in approved facilities) was granted by the Flemish administration (Agentschap voor Natuur en Bos, ID numbers ANB/BL-FFN 08-11344 and ANB/BL-FFN 12-00381). Behavioural assays were approved by the ethical committee of the University of Antwerp (ECD-dossier 2016-86, license approved 22-12-2016). Neither procedure adversely affected the starlings in the short term or for the overall period of the study. After each test session (see below), birds were returned to their holding conditions in their original large aviaries.

### Birds, housing and test room

The study was conducted on a captive population of adult starlings between January and April 2017 housed at the University of Antwerp campus in Wilrijk, Belgium. Only females were used to avoid seasonal effects, expected to be more marked in males. Formation of unisex male flocks could lead to aggression episodes already in early spring, while mixed-sex groups would generate behaviours linked to reproduction (Eens, 1997; Feare, 1984).

We used 56 adult females from two batches. The birds of the first batch (*n* = 24) were caught from the wild at several sites around Antwerp between 2005 and 2010. The birds of the second batch (*n* = 32) were also caught from the wild in the same sites in March 2016. From then onwards, they were held captive under the same standardized conditions. Female age (yearling or older) was estimated by measuring the mean tip length of three throat feathers (Komdeur et al., 2005). The first batch was composed of older females, and the second of yearling females. The first batch had been used for individual personality tests in 2016. Because of age differences as well as familiarity between individuals of the same batch, the two batches were always housed and treated separately during the different phases of the study. All birds were ringed with a numbered metal ring and a unique combination of plastic colour rings, allowing easy identification.

The birds of the two batches were housed into two large adjacent outdoor aviaries (each 6.0 × 16.0 × 3.0 m), each containing many perches. The food (Orlux® pate mixed with granule) was *ad libitum* available in a feeder, and *ad libitum* water was available in large trays and in a smaller bowl to allow drinking and bathing.

For both the individual test and the flock test (see below) we used the “exploration room”, a wooden structure (2.95 × 2.0 × 2.5 m) with a closed roof, three blind walls and a grid mesh in the front wall. This structure had already been used to quantify exploration in both male and female starlings (Fig. 1 in Thys et al., 2017a). The exploration room is a modified version of the open field test commonly used in avian personality studies (e.g., Carere et al., 2005; Dingemanse et al., 2002; Thys et al., 2017a; Verbeek et al., 1999).

**Figure 1.**
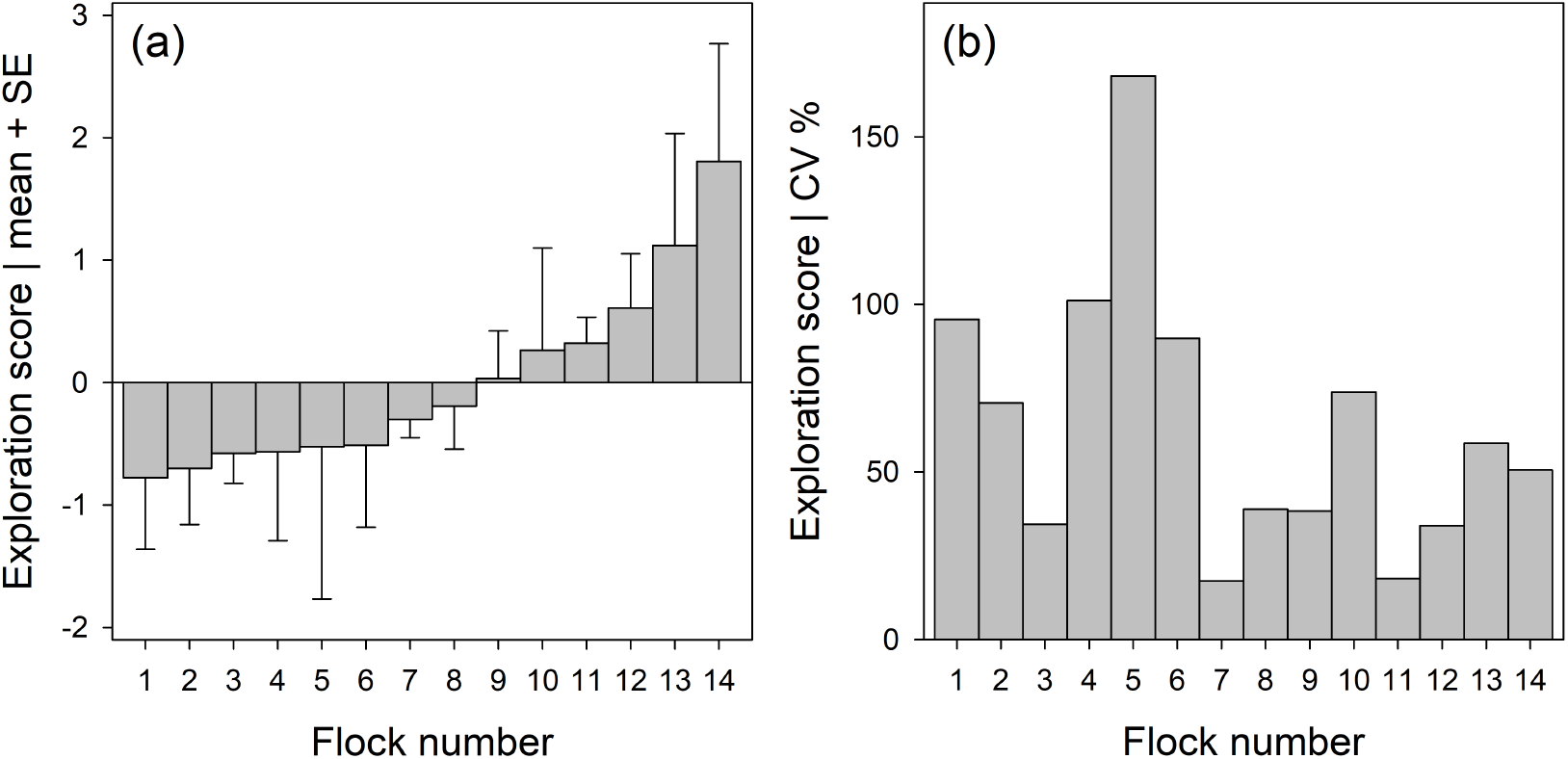
(a) Mean exploration score (+SE) and (b) the coefficient of variation (in %) within the different flocks, based on the individual exploration scores of the 4 individuals per flock. Flocks were designed after individuals were phenotyped by an individual-level exploration test.

Nine different items were present in the room: five perches, one platform, one “hole” (this is another entry of a start cage but closed), the grid and the ground. Birds could only change items by flying between them. Inside the test room, the temperature was similar to the external condition and true-light lamps with a flicker frequency of 100 Hz (fluorescent lamps that match closely the entire spectrum of natural day-light) were used.

Before the individual tests and during the whole part of the experiments (in which the birds were kept in flocks of 4 individuals) the birds were housed in 8 outdoor aviaries (3.2 × 2.0 × 2.5 m) with sand on the ground, and two perches over the width of the aviary. The aviaries were visually but not acoustically separated from each other.

### Individual tests

The tests were carried out in January 2017. Birds were caught from the big aviaries by two persons, and were moved with transportation cages into 7 of the 8 outdoor aviaries previously described, where they were housed in flocks of 7-10 females in each of them. After five days of habituation, 7-10 birds per day were each moved in individual cages (50 × 15 × 20 cm) with food and water. These cages were all located together in the one aviary that was purposefully kept empty. This moving was done between 4 and 5 pm to minimize stress and standardize the manipulation before the individual tests, which were carried out on the following day.

The following morning, each starling was transported in a cotton bag from the individual cage to the exploration test room (less than 1 min walking). The test was done during the time of the day when the birds are most active (between 8 am and 12 am). Observations were made from a hidden place attached to the room by a viewing window. All tests were video-recorded with Sony® Handycam HDR-XR550E/XR550VE, and the videos were scored with the software BORIS v3.43 (Friard & Gamba, 2016). The test started immediately after putting the bird in the “start cage” (24 × 14 × 14 cm) connected to the test room via an entry hole (5 cm diameter) at a height of 1.6 m, allowing birds to enter the room without further handling. A small sliding door giving access to the test room was lifted. The time taken for a bird to enter the room (latency to enter) was recorded. If a bird did not enter spontaneously the room after 1 min, it was stimulated by gently knocking the start cage. These birds (*n* = 5) had 60 seconds added to their latency. Behaviours were measured for 10 min after the bird had entered the room. For the first batch, as the same individuals had been tested the previous year (2016) in the same conditions, the data were used for the assessment of long-term repeatability. This allowed confirming the consistency of a personality trait in starlings (Thys et al., 2017a).

### Individual exploration score

We considered 4 measures both in 2016 (first batch) and 2017 (second batch) (and on same birds for the first batch): 1) latency to enter in the room, 2) total time flying (seconds), 3) total time spent on perches, 4) number of artificial trees visited. Based on these variables, a Principal Component Analysis (PCA) was run for each of the two batches, allowing to assign for each individual, a personality score based on the first principal component (see below). According to this score, we obtained a gradient of individuals, with slow and fast individuals at the extremes.

### Formation of flocks

We split the birds into groups of 4 according to their ranking and we formed 14 groups (6 groups for the first batch and 8 for the second batch) composed of individuals having all low exploration score or all high exploration score and all the intermediate levels (Fig. 1a). In doing so, we obtained groups with a gradient of individual personalities and also with a different coefficient of variation in individual exploration score (Fig. 1b). See details below, in the section on statistical analyses.

### Flock tests: exploration and perturbation test

#### Exploration test

This group-level test was based on the individual test with slight modifications: a larger start cage (25 × 25 × 27 cm), to contain 4 birds, and acclimatization time of 2 min. This was added to avoid any order effect due to the insertion of the birds in the cage one by one.

Similar to the individual test, the day prior the flock test, between 4 and 5 pm, 12 birds were moved into individual cages (we tested 3 flocks of 4 birds each per day). The next day (between 8 and 12 am), the birds were moved from their individual cage to the start cage and transported to the exploration room. During the test, the same 4 measures as in the individual exploration test were recorded. This allowed comparing the performance of each individual within the group to its performance when alone.

#### Perturbation test

After the standard 10 min of the group exploration test, the door of the small room in front of the exploration room where the birds were located, was suddenly opened and then closed creating a visual and acoustic perturbation for the birds. The birds were observed for 2 min after closing this ‘perturbation door’. We recorded the latency to fly after the perturbation and the latency to stop on an item after flying (see Suppl. Materials, Tables A-C for details).

### Statistical analysis and sample sizes

Analyses were based on two batches of animals (*n*_1_ = 24, *n*_2_ = 32) resulting in a total sample size of *n* = 56. All statistical analyses were done with R, version 4.2 (R Core Team, 2022).

First, we ran a Principal Component Analysis (PCA) across 3 repeated measurements of 4 behavioural variables (see above for details) observed in the animals of batch 1 (*n*_1_ = 24), twice in 2016 and once in 2017. We then checked for individual-based repeatability (*R*_ICC_) across time of the first Principal Component (PC), which was the only one with an eigenvalue >1. This was done by an intra-class correlation using a linear mixed-effects model (LMM) implemented in the R package *rptR* (Stoffel et al., 2017). *P*-values were calculated by Monte-Carlo sampling with 10,000 permutations.

Second, we ran another PCA based on the larger sample size (*n* = 56), as we integrated the additional 32 animals which we had behaviourally phenotyped only once in 2017. Again, we only used the first PC as exploration score, since all remaining components had eigenvalues < 1.

As next step, we checked whether relative within-group differences (calculated as the deviation from the group mean) in exploration score were associated with relative within-flock differences in the behaviours assessed during the exploration test and the perturbation test (*n* = 56 individuals). This was done by linear mixed-effects models (LMM) based on restricted maximum likelihood estimates, including group identity as a random factor using the *lmer* function of the R package *lme4* (Bates et al., 2015). For all LMM, we assessed the explained variation of significant associations by marginal *R*^2^ (Stoffel et al., 2021). Associations at the group level (*n* = 14) were calculated using linear models (LM). *P*-values for LMM were calculated using the *permlmer* function of the R package *predictmeans* (Luo et al., 2021), and *P*-values for LM were calculated by the *PermTest* function of the R package *pgirmess* (Giraudoux, 2016), always using 10,000 permutations. Permutation tests for linear (mixed-effects) models are well adjusted for moderate sample sizes and do not require normal distribution of model residuals (Good, 2005). However, as the error rate of permutation tests can be inflated when variances are unequal (Huang et al., 2006), we always checked for homogeneity of variances by plotting residuals versus fitted values (Faraway, 2006). The condition of variance homogeneity was obtained by log[x] transforming the latencies measured in the exploration and perturbation test, which originally showed a strong right-skewed distribution. This was also done for the following analysis.

As final step, we calculated different linear models explaining the birds’ flock-level responses (*n* = 14; exploration test, perturbation test) by different measures capturing between-flock differences in exploration tendency of the birds that made up these flocks (see Tables A, B in Suppl. Materials). Therefore, we calculated the mean, minimum and maximum individual exploration score within each flock, and also the coefficient of variation (CV), calculated as the standard deviation over the mean value. As the calculation of CV gives non-sensical results when based on positive and negative values (as it was the case for our PCA-based exploration score, ranging from approx. −2 to +3), we positivised these prior to CV calculation by adding a small, constant value (+2). The four measures obtained by our calculations of exploration scores within flocks (mean, min., max., CV) were not independent from each other, and thus we run different models and we compared these using AIC scores. We used the second order AIC, the AICc, which is well adjusted for comparisons based on smaller sample sizes (Burnham & Anderson 2002). All models of a ‘set’, i.e. with the same dependent variable (see Tables A-C in Suppl. Materials) were ordered from ‘best’ (lowest AICc) to ‘worst’ (highest AICc). By using AICc differences (ΔAICc = AICc*i* − AICc_minimum_) we compared the support that the different models found by the data. It has been suggested by Burnham and Anderson (2002) that models with ΔAICc*i* < 2 can be considered to have substantial support. Models with a ΔAICc of about 4 to 7 have considerably less support. We also calculated normalized Akaike weights (*w*i) for each model, which can be interpreted as a measure of the evidence that the model *i* is the best approximating model compared to the other models of the set (Burnham & Anderson, 2002).

## RESULTS

### Consistent individual differences in exploration tendency

We carried out a Principal Component Analysis (PCA) based on individual-level exploration tests, i.e., 3 repeated measurements from 24 individuals, which were tested twice in 2016 and once in 2017. During these tests, 4 different behavioural variables were quantified. The first Principal Component (PC1) explained 51.1% of the initial variance of the data, and the resulting PC1 score was positively associated with the time spent flying (loading: +0.598) and the number of artificial trees visited (+0.618), and was negatively associated with the latency to start exploring (−0.329) and with the time the individuals spent on perches (−0.390). This PC1 had an eigenvalue of 1.43; the eigenvalues of all remaining axes were < 1 and thus these axes were discarded for further analysis.

This PC1 score was significantly repeatable at the individual level between both test trials carried out in 2016 (LMM-based intra-class correlation with 10,000 permutations: *R*_ICC_ = 0.799, *n*_individuals_ = 24, *P* < 0.001), and between the first (*R*_ICC_ = 0.636, *n*_individuals_ = 24, *P* < 0.001) or the second test in 2016 (*R*_ICC_ = 0.599, *n*_individuals_ = 246., *P* < 0.001) and the test trial carried out in 2017 (see Fig. A, in Suppl. Materials).

For the following analyses, we calculated a PCA using the same input variables only for the test trial in 2017, based on a larger sample size (*n*_individuals_ = 56) containing 32 additional individuals for which we only had taken a single measure of individual exploration. In accordance with the results described above, the first PC again explained a large part (47.0%) of the total variance of the data and the loading showed the same signs, respectively: the time spent flying (+0.655), the number of artificial trees visited (+0.656), the latency to start exploring (−0.151) and the time the individuals spent on perches (−0.342). The PC1 scores did not significantly differ between animals from the two batches (LM with 10,000 permutations: *P* = 0.121).

### Formation of flocks

Based on the PC1 score (exploration score) obtained for each bird (*N* = 56), we designed 14 flocks of 4 individuals each. These groups were designed in such a way that a gradient of mean exploration scores per flock emerged (Fig. 1a), ranging from −0.78 ± 0.58 SE (low average exploration tendency) to 1.81 ± 0.96 SE (high average exploration tendency). Furthermore, the flocks differed in the coefficient of variation in individual exploration scores, ranging from 17.53% to 168.19% (Fig. 1b).

There was no significant correlation between the coefficient of variation and the average exploration score per flock (LM with 10,000 permutations: *R*^2^ = 0.128, *β* = 0.358 ± 0.269 SE, *P* = 0.208).

### 3.3 Individual exploration score and individual behaviour in flocks

Our results indicate that individuals with a relatively higher individual exploration score compared to their group mates (calculated as the deviation from the group mean) showed relatively shorter latencies in a flock-level exploration test and perturbation test. This was supported by significant and negative associations between the relative individual exploration score and the relative latencies to to enter the arena during the exploration test (LMM with 10,000 permutations: *marginalR*^*2*^ = 0.071, *β* = −0.269 ± 0.131 SE, *P* = 0.046; Fig. 2a) and to stop flying after the perturbation (*marginalR*^*2*^ = 0.100, *β* = −0.319 ± 0.129 SE, *P* = 0.021; Fig. 2c), and by the non-significant tendency of a negative association between the relative individual exploration score and the relative individual latency to start flying in the perturbation test (*marginalR*^*2*^ = 0.069, *β* = −0.263 ± 0.131 SE, *P* = 0.057; Fig. 2b).

**Figure 2.**
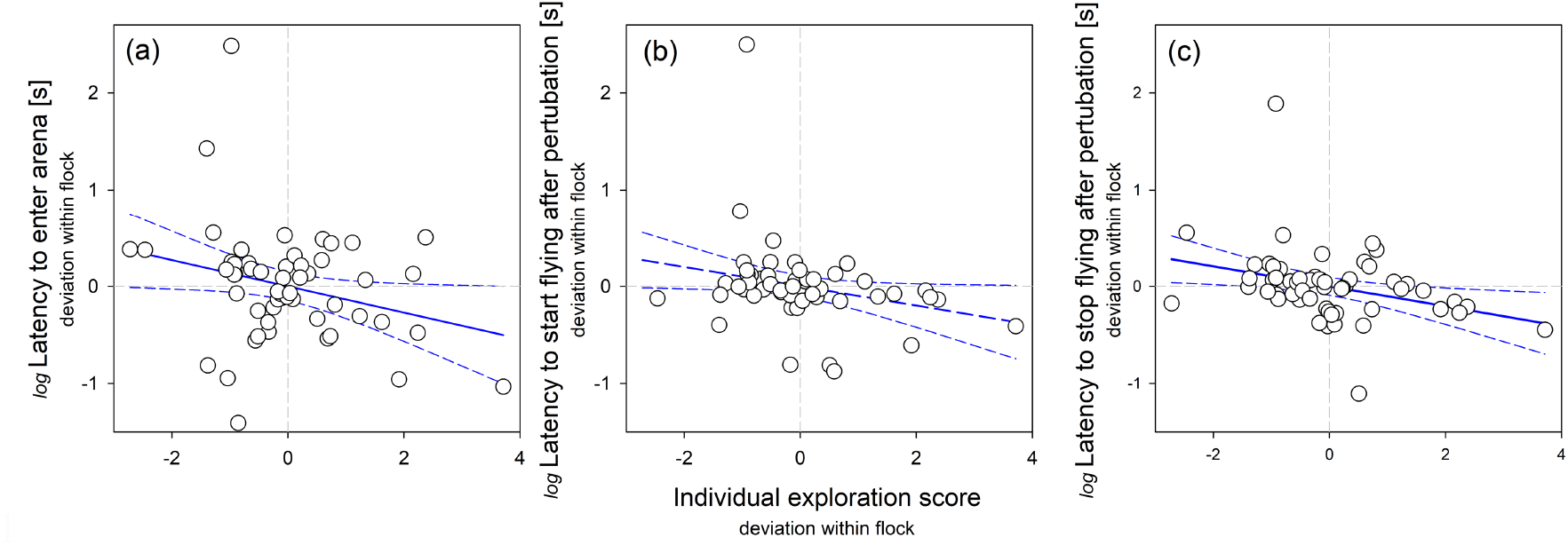
Associations between the individual exploration score and the within-flock differences in individual (log-transformed) latencies (a) to enter the test area during the exploration test, and (b) to start flying as well as (c) to stop flying after a perturbation. Associations shown in (a, c) were statistically significant and the association in (b) showed a statistical tendency; see text for details. 95% confidence intervals around the regression lines are given as dashed lines. Analyses were based on a total of 56 individuals out of which 14 flocks were designed (4 individuals each).

### 3.4 Variation of individual exploration scores within the flock and flock-level behaviour

#### Exploration test

the within-flock variability in individual exploration scores (as assessed by a coefficient of variation) was significantly and positively correlated with the range span in latency (LM with 10,000 permutations: *R*^2^ = 0.395, *β* = 0.629 ± 0.225 SE, *P* = 0.029; Fig. 3a) as well as with the mean latency (*R*^2^ = 0.373, *β* = 0.619 ± 0.228 SE, *P* = 0.020; Fig. 3b) until the birds of a flock had entered the test arena during the exploration test. That is, flocks consisting of individuals with a more homogenous distribution of exploration scores entered the test arena within a shorter span of time and on average more quickly than more heterogeneous flocks. Furthermore, there was also a significant and positive correlation between the minimum individual exploration score occurring in the flock and the mean flock latency to enter the test arena (*R*^2^ = 0.309, *β* = 0.556 ± 0.240 SE, *P* = 0.040). A comparison of these two competing models, which explained variation in the mean flock latency either by the coefficient of variation or by the minimum individual exploration per flock by their AICc scores indicated that the former model found slightly better support by the data (Δ*AICc* = 1.363; see details in Table A in the Suppl. Materials). All other models explaining the range span or the mean flock latency by the mean or maximum individual exploration score of the flock members were not statistically significant and had a considerably lower AICc score (see Table A in Suppl. Materials).

**Figure 3.**
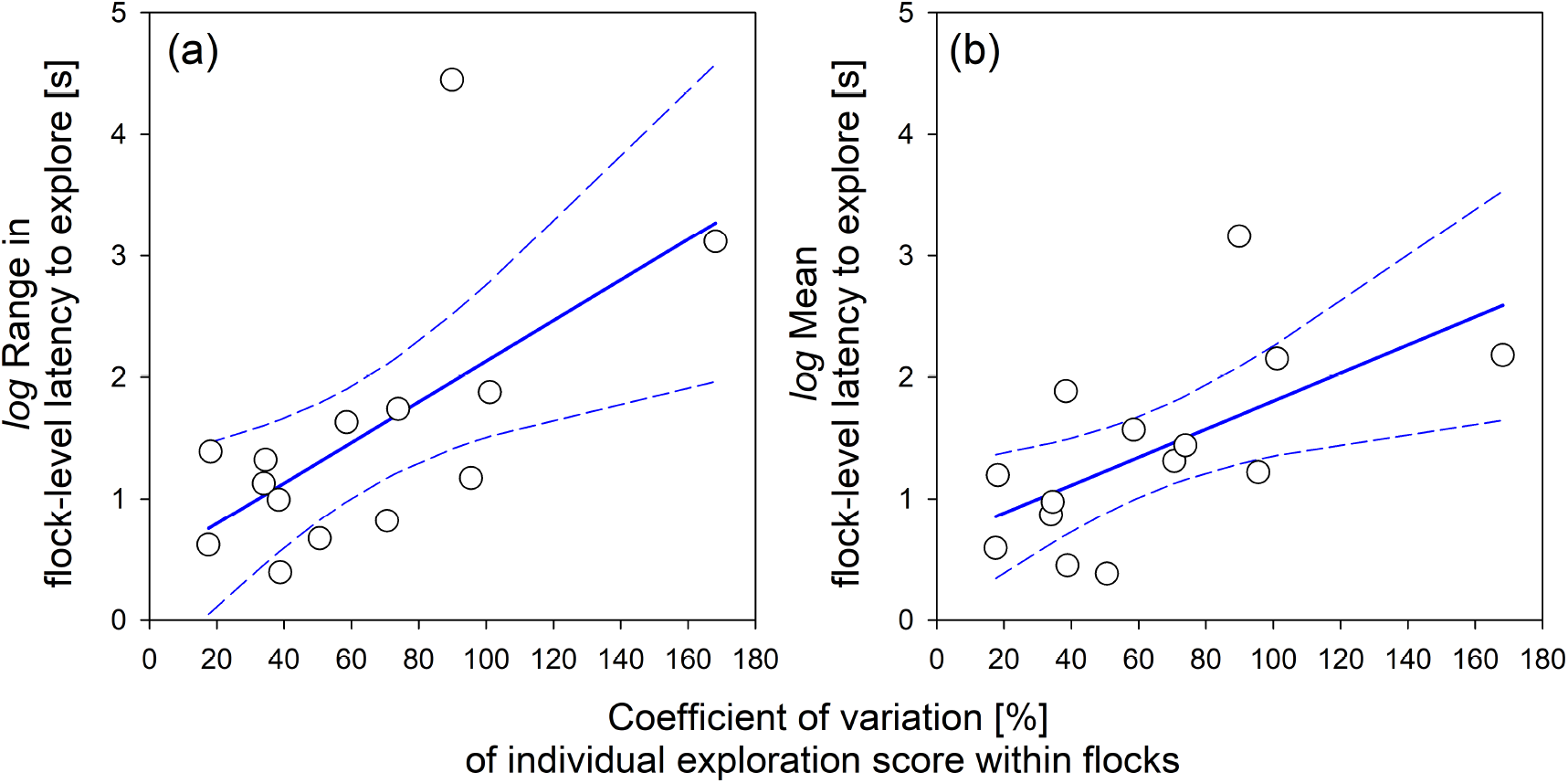
Flock-level associations between the coefficient of variation of the individual exploration score within groups and (a) the range (maximum-minimum) or (b) the mean latency until the birds (*n*_individuals_ = 56) of the different flocks (*n*_flocks_ = 14) had entered the test arena during the exploration test. 95% confidence intervals around the regression lines are given as dashed lines. Both regressions were statistically significant, see text for details.

#### Perturbation test

we also tested for correlations between the coefficient of variation, the average, and the maximum and minimum individual exploration score and the flocks’ responses in a perturbation test, i.e., the average and the range span in latency to start (escape response) and to stop flying activity. However, none of these correlations were statistically significant (all *P* > 0.10; see details in Tables B, C in Suppl. Materials).

There were no significant differences in any of the tested flock-level behaviours for between flocks composed of animals from the two different batches, neither during the exploration test, nor during the perturbation test (all *P* > 0.05).

## DISCUSSION

Theoretical and empirical studies investigating the behavioural phenotype of social groups taking into account the role of the personality of the individual composing them have recently increased (e.g., Cook et al., 2022; Carere et al., 2018; Herbert-Read, 2017; Munson et al., 2021; Pinter-Wollmann, 2012; Vágási et al., 2021;), but almost nothing is known about the effect of within-group homogeneity in personality traits on group behaviour.

In this study, we investigated the impact of consistent individual differences in a personality trait on group responses by experimentally creating flocks of starlings with different degrees of intra-flock variation. We focussed in particular on the functional role of within-group phenotypic homogeneity, since so far this group attribute has been not studied concerning behavioural traits reflecting personality, such as exploration (see introduction), despite there has been an emphasis on highlighting the possible advantages of group diversity in personality. Here, we tested whether different degrees of within flock individual variation in exploratory behaviour affect flock performance in the same task and in escape response upon a frightening stimulus.

Our study first confirmed previous findings reporting long- and short-term individual behavioural consistency across time in the European starling when tested individually for exploration (Thys et al., 2017a, b). On the basis of these individual exploration scores, we then formed a “gradient” of same-sized flocks differing in both mean and internal variation of exploration scores, thereby in within-flock homogeneity. Later on, the behaviour of the individuals in the flocks was scored in two group tasks, an exploration and a perturbation test. At the individual level, in these group tasks our results indicate that birds with a relatively higher individual exploration score compared to their group mates (calculated as the deviation from the group mean) showed, or tended to show, relatively shorter latencies. At the group level, flocks composed by individuals with a more homogenous distribution of exploration scores entered the test arena within a shorter span of time and on average more quickly than more heterogeneous flocks. No significant effects on escape response to a perturbation emerged.

The social conformity hypothesis predicts that individuals in a group can either synchronize their behaviour towards the ‘middle’ or change it towards that of certain members, and this should improve group performance (Brown & Irving, 2014; Fürtbauer & Fry, 2018; Jolles et al., 2017). However, if the starting point is already a high degree of phenotypic homogeneity, the conformity could be achieved more easily and rapidly. This is likely in the case of highly gregarious species (bird flocks and fish schools) with fission-fusion dynamics where group composition changes frequently (Couzin & Laidre, 2009) in association with complex collective behaviours that include high frequencies of mixing, merging and splitting as in the case of starlings (Storms et al., 2019). Since these flocks base their collective behaviour on self-organization rules (Hildebrandt et al., 2010), it could be hypothesised that in nature starlings might associate with the flock that maximises phenotypic homogeneity in personality traits. This hypothesis, not tested so far, entails the assumption that individuals are somehow able to perceive/assess the personality of the other members of a flock when deciding to join it or not. Such assessment could occur at the typical social roosts that this species forms, especially during autumn/winter (Carere et al., 2009; Feare 1984). An interesting perspective for future experiments in controlled conditions could be to explore individual associations while roosting and sleeping in relation to personality traits.

We did not find any significant effects of flock homogeneity on the second group parameter tested, the escape response to a perturbation test measured as latency to start again flying activity and to stop on an item after flying. A similar sudden ‘disturbance’ test has been applied to small groups of ants composed by individuals with different exploratory tendencies. The disturbance consisted in lifting and agitating the nest tube so that the individuals and the brood (cocoons) dropped; then, the ants were offered a new nest tube and the number of cocoons as well as the latency to bring the cocoons back to the nest, the number of individuals going to the nest, and their latency were recorded (Carere et al., 2018). Groups with more exploratory individuals started transporting displaced cocoons significantly earlier and transported more cocoons into the nest than groups with low exploratory individuals.

In the case of starling flocks, the lack of group effect following a perturbation may simply be due to the fact that, while exploration is a spontaneous activity entailing lower potential risks, a sudden perturbation also fostered by visual risk assessment processes, well known in starlings (e.g., Devereux et al., 2006), means danger. If there are no predators or other risks around, the individual would follow the behaviour of the flock. Conversely, in case of an actual sudden danger that is similarly perceived by all flock members as in our perturbation test, the behaviour of flock mates is, at least initially, disregarded. This could be comparable to the common aerial response that the starling flocks immediately display in case of a predator attack at high speed, the so-called ‘flash expansion’. This collective escape response, also common in insects and fish, is characterised by individuals suddenly moving radially outward the group, which loses cohesion (Romey & Lamb, 2015; Storms et al., 2019).

In conclusion, our study highlights the relevance of incorporating not only consistent individual variation, but also the within-group homogeneity of consistent behavioural types in models of social/collective behaviour.

## Author Contributions

C.C. and P.dE conceived the study, planned the experimental design and methodology and acquired funding. C.C., P.dE and H.G.R. wrote the paper. H.G.R analysed the data. M.E. and R.P. provided the facilities, the animals, and contributed to experimental set-ups and manuscript revisions. C.C., C.A. and F.D. carried out the experiments. C.C. supervised the students. All authors reviewed and commented on drafts of the manuscript and gave final approval for publication.

## Declaration of interest

The authors declare they have no competing interests.

## Acknowledgments

We are very grateful to Geert Eens and Peter Scheys for excellent animal care and support during the experiments. The study was financially supported by the Marie Sklodowska-Curie Action H2020-MSCA-IF-2014-659106 (GROUPIND). P.dE. is funded by a grant from the Institut Universitaire de France (IUF). The study was part of the Master’s theses of C.A. (Ethology) and F.D. (Applied Ethology) at the Université Sorbonne Paris Nord.

## Supplementary Materials

**Figure A.**
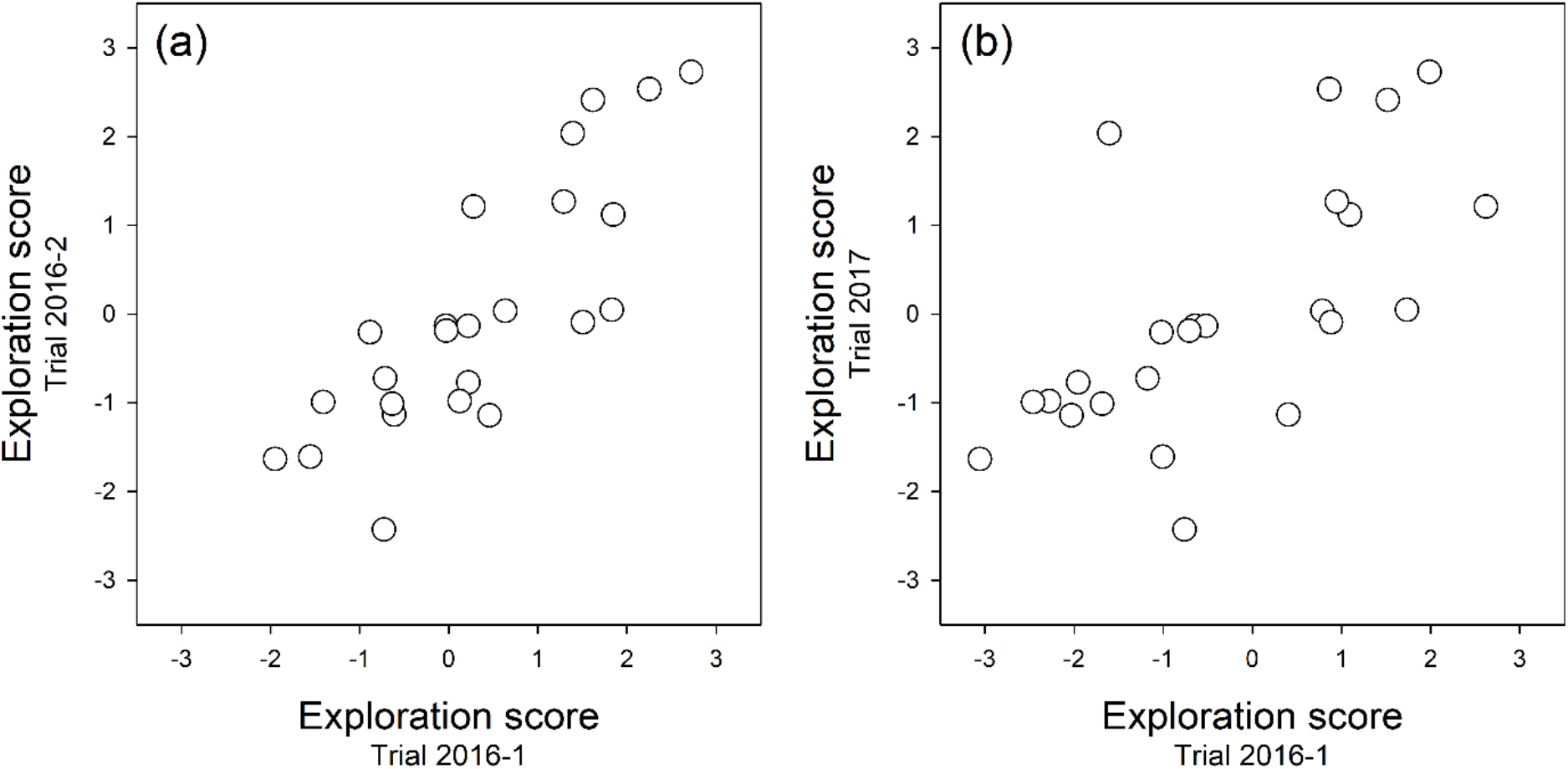
Individual-level consistencies across time in exploration scores, as obtained by the first axis of a PCA based on the animals’ latency to enter the test arena, their time spent flying, the number of trees visited and the time spent sitting on a perch. Animals (*n* = 24) were repeatedly tested: twice in 2016 (with an interval of 2 weeks) and once in 2017. Intra-class correlations of the data shown in (a), short-term repeatability, and in (b), long-term repeatability, were significant (see main text for details).

**Table A.**
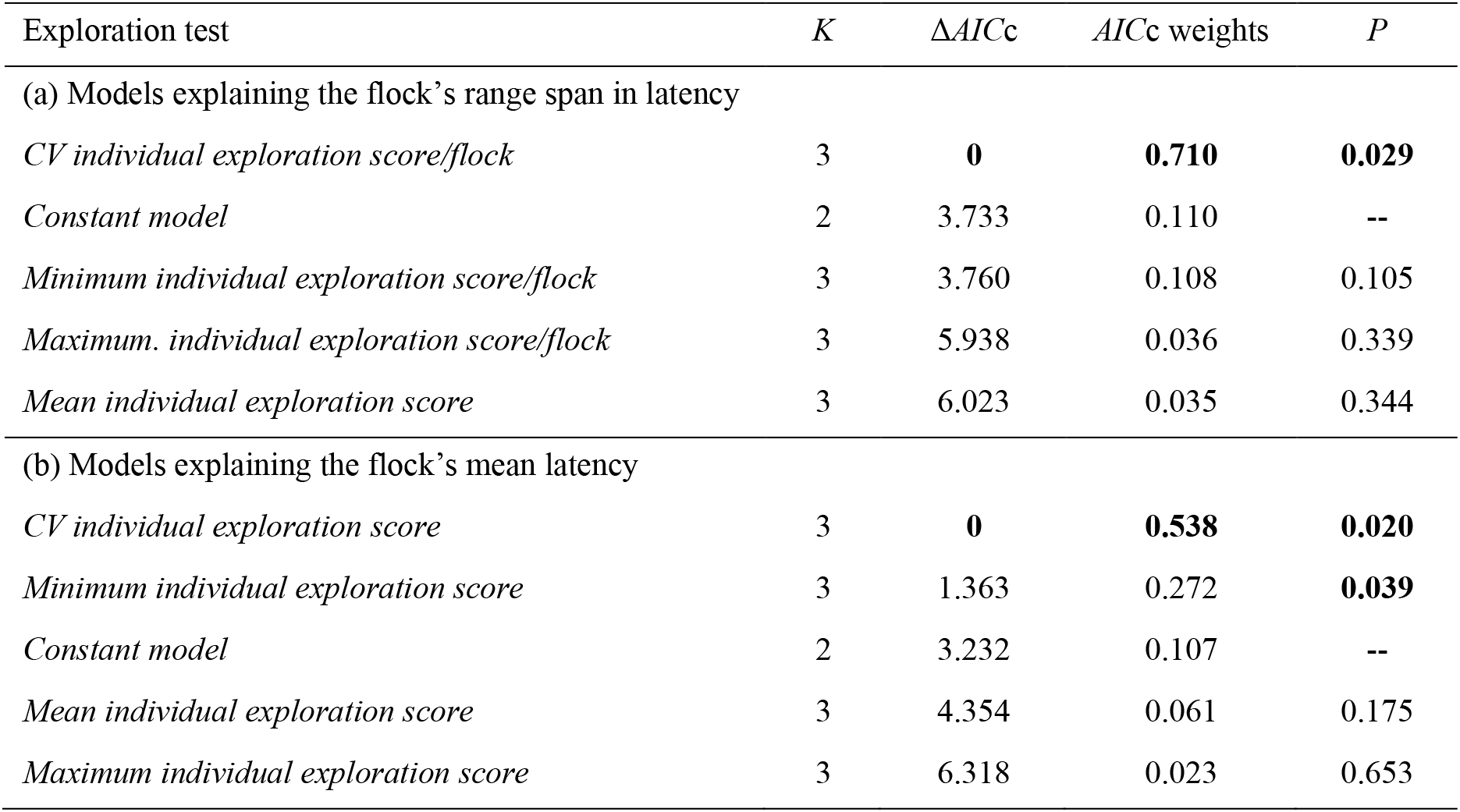
Model sets (linear models) explaining the animals’ responses in a **flock-level exploration test** by different measures capturing the distribution of individual differences in exploration tendency (exploration score) within flocks; the latter was previously determined in individual-based tests. Two different (log-transformed) flock-level dependent variables were considered: (a) the range (maximum-minimum) and (b) the average latency until the birds of *n* = 14 different flocks (4 animals per flock) had entered the test arena. The number of estimable parameters (*K*), the relative differences of Akaike’s information criterion (Δ *AIC*c) corrected for small sample sizes, Akaike weights, and *P*-values are given for each model within each set (a, b). *P*-values were calculated by 10,000 Monte Carlo permutations.

**Table B.** Model sets (linear models) explaining the animals’ responses in a **flock-level perturbation test** by different measures capturing the distribution of individual differences in exploration tendency (exploration score) within flocks; the latter was previously determined in individual-based tests. Two different (log-transformed) flock-level dependent variables were considered: (a) the range (maximum-minimum) and (b) the average latency until the birds of *n* = 14 different flocks (4 animals per flock) **had started to fly after the perturbation**. The number of estimable parameters (*K*), the relative differences of Akaike’s information criterion (Δ*AIC*c) corrected for small sample sizes, Akaike weights, and *P*-values are given for each model within each set (a, b). *P*-values were calculated by 10,000 Monte Carlo permutations.

| Perturbation test | $K$ | $\Delta AICc$ | $AICc$ weights | $P$ |
| --- | --- | --- | --- | --- |
| (a) Models explaining the flock's range span in latency |  |  |  |  |
| *Constant model* | 2 | **0** | **0.513** | -- |
| *Mean individual exploration score/flock* | 3 | 2.587 | 0.141 | 0.436 |
| *Maximum individual exploration score/flock* | 3 | 2.832 | 0.125 | 0.548 |
| *Minimum individual exploration score/flock* | 3 | 2.909 | 0.120 | 0.599 |
| *CV individual exploration score/flock* | 3 | 3.247 | 0.101 | 0.816 |
| (b) Models explaining the flock's mean latency |  |  |  |  |
| *Constant model* | 2 | **0** | **0.538** | -- |
| *Mean individual exploration score/flock* | 3 | 2.900 | 0.126 | 0.543 |
| *Minimum individual exploration score/flock* | 3 | 2.985 | 0.121 | 0.616 |
| *CV individual exploration score/flock* | 3 | 3.218 | 0.108 | 0.767 |
| *Maximum. individual exploration score/flock* | 3 | 3.221 | 0.107 | 0.750 |

**Table C.** Model sets (linear models) explaining the animals’ responses in a **flock-level perturbation test** by different measures capturing the distribution of individual differences in exploration tendency (exploration score) within flocks; the latter was previously determined in individual-based tests. Two different (log-transformed) flock-level dependent variables were considered: (a) the range (maximum-minimum) and (b) the average latency until the birds of *n* = 14 different flocks (4 animals per flock) **had stopped flying around after being startled by the perturbation**. The number of estimable parameters (K), the relative differences of Akaike’s information criterion (delta *AIC*c) corrected for small sample sizes, Akaike weights, and *P*-values are given for each model within each set (a, b). *P*-values were calculated by 10,000 Monte Carlo permutations.

| Perturbation test | $K$ | $\Delta AICc$ | $AICc$ weights | $P$ |
| --- | --- | --- | --- | --- |
| (a) Models explaining the flock's range span in latency |  |  |  |  |
| *Constant model* | 2 | **0** | **0.548** | -- |
| *Maximum. individual exploration score/flock* | 3 | 3.069 | 0.118 | 0.638 |
| *CV individual exploration score/flock* | 3 | 3.083 | 0.117 | 0.654 |
| *Mean individual exploration score/flock* | 3 | 3.248 | 0.108 | 0.827 |
| *Minimum individual exploration score/flock* | 3 | 3.249 | 0.108 | 0.824 |
| (b) Models explaining the flock's mean latency |  |  |  |  |
| *Constant model* | 2 | **0** | **0.521** | -- |
| *Maximum. individual exploration score/flock* | 3 | 2.384 | 0.158 | 0.372 |
| *CV individual exploration score/flock* | 3 | 3.019 | 0.115 | 0.619 |
| *Mean individual exploration score/flock* | 3 | 3.178 | 0.106 | 0.730 |
| *Minimum individual exploration score/flock* | 3 | 3.305 | 0.100 | 0.952 |

## Notes

### Competing Interest Statement

The authors have declared no competing interest.

